## Supplementary material for "Pragmatic representations of self- and others’ action in the monkey putamen": Supplementary materials.pdf

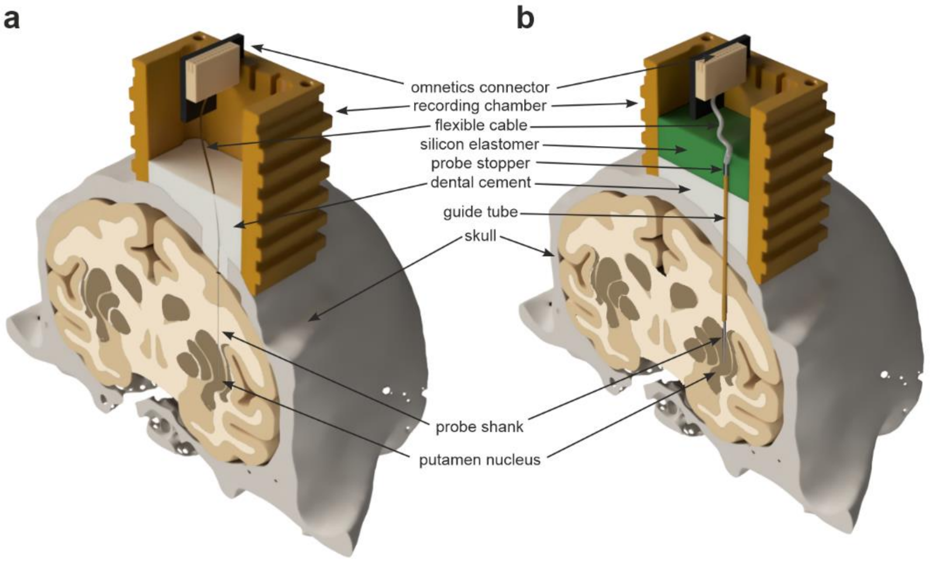


**Supplementary Fig. 1 | Schematic of silicon and stainless-steel probe implants**. **a**, Chronic implant of silicon probes in Mk1. The trajectory deviation caused by the probe's angled tip was measured in vitro using agarose, revealing a deflection of 0.05 to 0.12 mm per millimeter of insertion. Over the full 24 mm insertion path, this resulted in a cumulative deviation of 1.2-2.9 mm toward the bevel-opposite side. This deviation was taken into account during the surgical insertion procedure to ensure accurate placement of the recording shaft. **b**, Semi-chronic implant in Mk2 using stainless-steel multisite linear probes inserted through permanently implanted Teflon guide tubes. Further details regarding chamber layout and data logging have been described elsewhere[60].

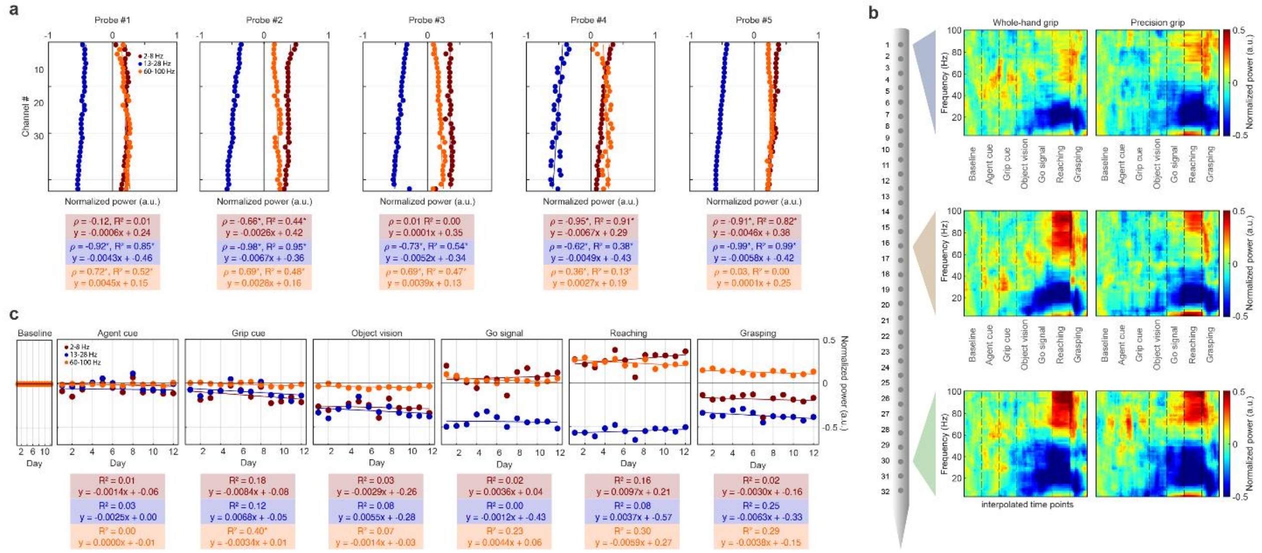


**Supplementary Fig. 2 | Depth- and time-dependent modulation of Local Field Potentials (LFP) power**. **a**, Depth-dependent modulation of LFP power across all channels. Each panel represents a separate probe. Regression lines, along with the corresponding Spearman correlation coefficients (ρ) and $R^2$ values, quantify the relationship between depth and LFP power for each frequency band. **b**, Representative examples of LFP activity across different depths, emphasizing a highly reproducible temporal modulation pattern in the investigated frequency bands across depths. **c**, Time-dependent modulation of LFP power across recording days. Power fluctuations in each frequency band are shown for different task epochs. Regression analyses reveal no significant drift in power across days, confirming stable recording conditions. * p<0.05.

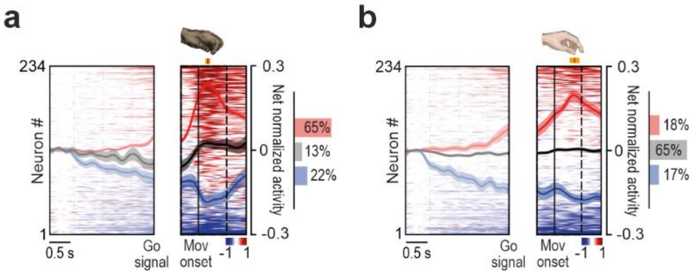


**Supplementary Fig. 3 | Neuronal properties of facilitated and suppressed task-related neurons during self and other trials**. **a**, Average population response overlapped on single neuron heatmaps ordered by the magnitude of response around the Movement onset [-0.3 to +0.5 s] during monkey's own action. Population activity of facilitated (red), suppressed (blue), and non-motor-related (black) neurons. **b**, Average population response overlapped on single neuron heatmaps ordered as in a) during experimenter's action. In both plots, orange markers indicate the average ± 1 standard deviation time at which the hand makes contact with the object. Histograms on the right of each plot illustrate the fraction of neurons exhibiting significant facilitation (red), suppression (blue), or non-significant modulation in that task condition.

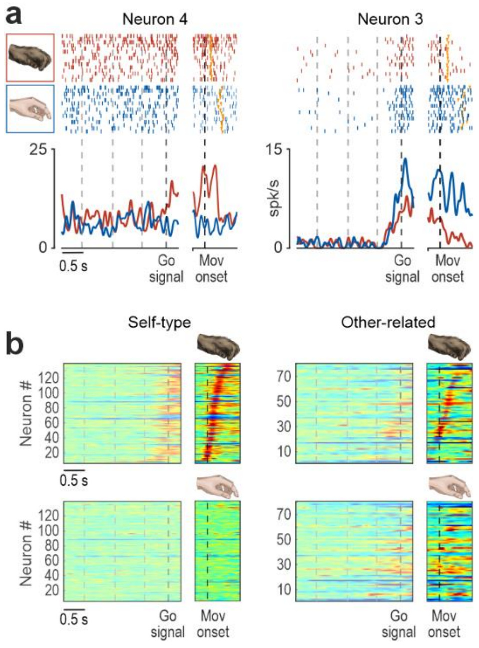


**Supplementary Fig. 4 | Simultaneously recorded neurons during self and other trials**. **a**, Examples of self-type (Neuron 4) and self-and-other-type (Neuron 3) cells simultaneously recorded during monkey's own (red) and experimenter's (blue) actions. Neuron 4 discharged during self-trials but remained silent in the same other-trials in which Neuron 3 was active. Conventions as in panel 4a. **b**, Activity heat maps of self-type and other-related neurons during the same monkey and experimenter trials. Alignments as in Fig. 4a. Neurons are ordered (bottom to top) according to the timing of their peak of activity during the monkey's own action execution. Note that neurons discharging during the monkey's active movement are silent during the experimenter's actions, when other-related neurons fire, suggesting that subtle monkey movements are unlikely to occur and hence can hardly explain the observed modulation of other-related neurons activity during the experimenter's action.

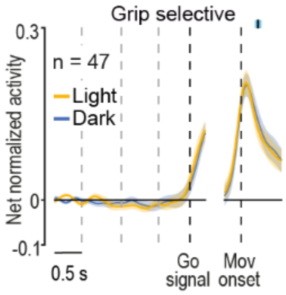


**Supplementary Fig. 5 | Population response of grip-selective neurons in their preferred grip condition during self-trials performed in the light and in the dark**.
